# Binding Entropy Can Be Predicted by Crystallographic Ensembles

**DOI:** 10.64898/2026.01.19.699807

**Authors:** Charlotte A. Miller, Stephanie A. Wankowicz

## Abstract

Protein-ligand binding is governed by free energy, comprising both enthalpic and entropic contributions. Yet structural interpretations of binding thermodynamics have predominantly focused on enthalpic interactions, largely neglecting entropy because it is difficult to quantify from static structural models. Here, we developed multiconformer ensemble models to analyze high-resolution X-ray crystallography structures and estimate both protein and solvent conformational entropies. These ensemble models successfully predicted experimental binding entropies measured by isothermal titration calorimetry for over 70 protein-ligand pairs across 12 proteins, revealing a strong linear correlation. Protein entropy, estimated using crystallographic order parameters that capture both harmonic and anharmonic motion, correlates linearly with experimental binding entropy. Incorporating resolution-corrected differences in water-molecule counts substantially improves predictions, demonstrating that protein and solvent contributions must be considered jointly. Analysis of water-protein hydrogen bonding networks partially explains entropic differences across complexes. These results establish that crystallographic ensembles can quantify binding entropy, enabling explicit entropic considerations in structure-based studies of molecular recognition for both functional analysis and drug design.

## Introduction

Protein–ligand interactions orchestrate enzyme function and signaling throughout biology. Much of modern medicine harnesses these interactions to manipulate proteins with synthetic ligands, thereby altering disease pathways. We have long sought to understand the physical principles underlying protein-ligand interactions. However, our static structural models, and thus our physical interpretations, have historically emphasized enthalpic interactions[1]. Yet binding affinity is determined by free energy, which comprises both enthalpic and entropic contributions. The entropic component of free energy has received far less attention, largely because entropy cannot be directly visualized or quantified from static structural models and instead requires ensemble representations [2,3]. Quantifying binding entropy directly from structural data would enable systematic entropic optimization and help reveal the structural determinants of binding thermodynamics.

Explanations of the entropic component of ligand binding often focus on solvent molecule release [4][5]. Yet accumulating evidence indicates that protein entropy is also a major determinant of binding entropy, with contributions differing substantially between proteins and between different ligands binding to the same protein [2,6–11]. Protein conformational entropy arises from the many degrees of freedom proteins possess, much of which is derived from side-chain sampling within and between rotamer states [12–15]. This conformational multiplicity, along with solvent entropy, creates an entropic reservoir that ligand binding perturbs, contributing to both protein stability and binding thermodynamics [6,12,16–21].

Developing quantitative metrics to estimate protein entropy has been challenging. Protein entropy encompasses both conformational entropy, which reflects anharmonic transitions between discrete rotamer states, and harmonic vibrational entropy, which reflects fluctuations within individual rotamer states. Accurate entropy calculations require enumerating the partition function over all conformational states, weighted by their Boltzmann factors [22]. However, this is computationally intractable due to the high dimensionality of conformational space, leading to various quantitative approximations. In molecular dynamics (MD), quasi-harmonic analyses of trajectories approximate entropy from covariance matrices but fail to capture the anharmonic entropic contributions [23,24]. Further, many MD simulations do not span biologically relevant timescales[21]. Nuclear Magnetic Resonance (NMR) relaxation techniques provide site-specific measurements of disorder in methyl or amide groups on picosecond timescales [25,26]. These measurements capture probability distributions of spatial coordinates and have been correlated with entropy [7,14,27]; but face challenges of throughput, timescale, and structural degeneracy.

X-ray crystallography and cryo-electron microscopy (cryo-EM) collect data from millions of protein copies, capturing many conformational states, therefore potentially containing significant information on the multiplicity of states and, thus, entropy. However, structural models typically represent only a single state, discarding information about the multiplicity of states that drive entropy [28]. We previously demonstrated algorithmic advances that enable modeling high-resolution X-ray crystallography data as multiconformer ensembles [28,29]. These ensembles explicitly capture anharmonic heterogeneity, including differences in rotamer states through alternative conformations, as well as harmonic heterogeneity through B-factors, thereby improving fit to experimental data [29]. Critically, these models enable quantitative estimates of protein conformational entropy via crystallographic order parameters [30], yielding values that agree with NMR relaxation order parameters and thus capture the spatial distribution of side chains.

Previous studies identified a quantifiable relationship between binding entropy and protein conformational entropy as measured by NMR order parameters [30,31], leading to the development of the ‘entropy meter’ [7]. Recently, we demonstrated that within a single system, protein and solvent entropy derived from crystallographic ensembles correlated with binding entropy [8]. We also showed that protein-solvent hydrogen-bonding networks correlate with binding entropy, linking protein conformational entropy to solvent entropy through coordinated, system-level redistribution. However, whether these findings generalize beyond individual systems remains an open question.

Here, we establish that crystallographic ensembles provide quantitative entropy estimates that correlate with binding entropy across multiple systems. We show that accounting for both protein and solvent entropy is key to predicting binding entropy. These structure-based entropy metrics enable integration of this critical free energy component into both the intuition and quantification of binding thermodynamics.

## Results

### Dataset

We collected thermodynamic binding measurements from isothermal titration calorimetry (ITC) from our previous study and from 14 datasets representing 12 different proteins (SARS-CoV-2 macrodomain [Mac1], Cyclin-dependent kinase 2 [CDK2], Thermolysin, Major Urinary Protein 1 [MUP-1], Bromodomain-containing protein 4 [BRD4], tRNA Guanine Transglycosylase [TGT], human Carbonic Anhydrase II [HCAII], Farnesyl pyrophosphate synthase [FPS], Galectin, and Transthyretin [TTR], Bovine Trypsin [Trypsin]), totaling 75 unique protein-ligand interactions(**Supplementary Table 1**) [8,9,32–41]. For protein-ligand pairs with multiple ITC measurements, we averaged each measurement for subsequent analysis. For each ITC measurement, we required a corresponding structure containing the same protein-ligand pair measured by ITC, along with an apo structure with the same sequence, space group, and resolution (within 0.3 Angstroms [Å]) and unit cell dimensions/angles (within 10%) (**Supplementary Table 2**). While we looked at additional protein-ligand pairs, most notably the extensive work on HIV protease and Thrombin [42–45], these did not meet the stringent crystallographic criteria (see all datasets examined but omitted in **Supplementary Table 3**). All structures had resolutions better than 2.1 Å, and their structure factors were deposited (**Supplementary Figure 1A/B**). We remodeled all structures using a pipeline of multiconformer models from qFit and Phenix refinement (**Methods**) [29,46,47]. We required multiconformer models to have an Rfree better than 0.25 (**Supplementary Figure 1C/D**). Three pairs were removed due to refinement errors.

Our dataset comprised proteins with bound chains ranging from 115 to 316 residues (median = 258; **Supplementary Figure 2A**). All bound and unbound structure pairs exhibited minimal conformational changes, with alpha-carbon root-mean-square deviations (RMSD) below 1.75 Å (**Supplementary Figure 2B**). This confirmed the absence of large conformational changes, as previously investigated by NMR [31]. Ligands ranged in molecular weight from 102.46 to 461.50 Da (median = 284.68), with hydrogen bond acceptor counts of 1–7 (median = 4) and hydrogen bond donor counts of 0–5 (median = 2; **Supplementary Figure 3A**).

### Binding Entropy Linearly Correlates with Protein Conformational Entropy

We first tested whether protein entropy, estimated from crystallographic ensembles, correlated with binding entropy (**Figure 1A**). To estimate protein entropy from multiconformer structures, we calculated crystallographic order parameters that measure the first torsion angle of each alternative conformation, weighted by its B-factor. We measured them for all residues except glycine and proline (**Methods**) [48]. We calculated the difference in protein entropy as the average difference between residues multiplied by the number of residues in the structure, as previously done [7]. When we compared these structural estimates to binding entropy measurements from ITC, we observed a linear relationship but found that our structural estimates systematically overestimated entropy changes by approximately fourfold. After applying a correction factor of 0.25, the structural estimates aligned much more closely with experimental values along a 1-to-1 line. Proteins showing increased conformational entropy in the bound state exhibited more entropically driven binding (Slope=0.85, R²=0.59; **Figure 1B; Supplementary Table 4/5**). We also explored the correlation between protein conformational entropy and individual measured ITC values and found a significant correlation, though with greater scatter around the median trend line, as expected given experimental variation in replicate measurements (**Supplementary Figure 4**). This correlation indicates that binding entropy increases by approximately 0.71kcal/mol·K per unit of protein conformational entropy. The magnitude of these changes provides insight into the distribution of protein structures. Given that the proteins examined average 258 residues and that the maximum conformational entropy change per residue is 0.25 kcal/mol·K (with a median closer to 0.05 kcal/mol·K), the observed entropy changes likely arise from tens to hundreds of residues shifting their conformational sampling. This result also indicates that changes in protein conformational entropy are a major component of the overall ligand-binding entropy.

**Figure 1.**
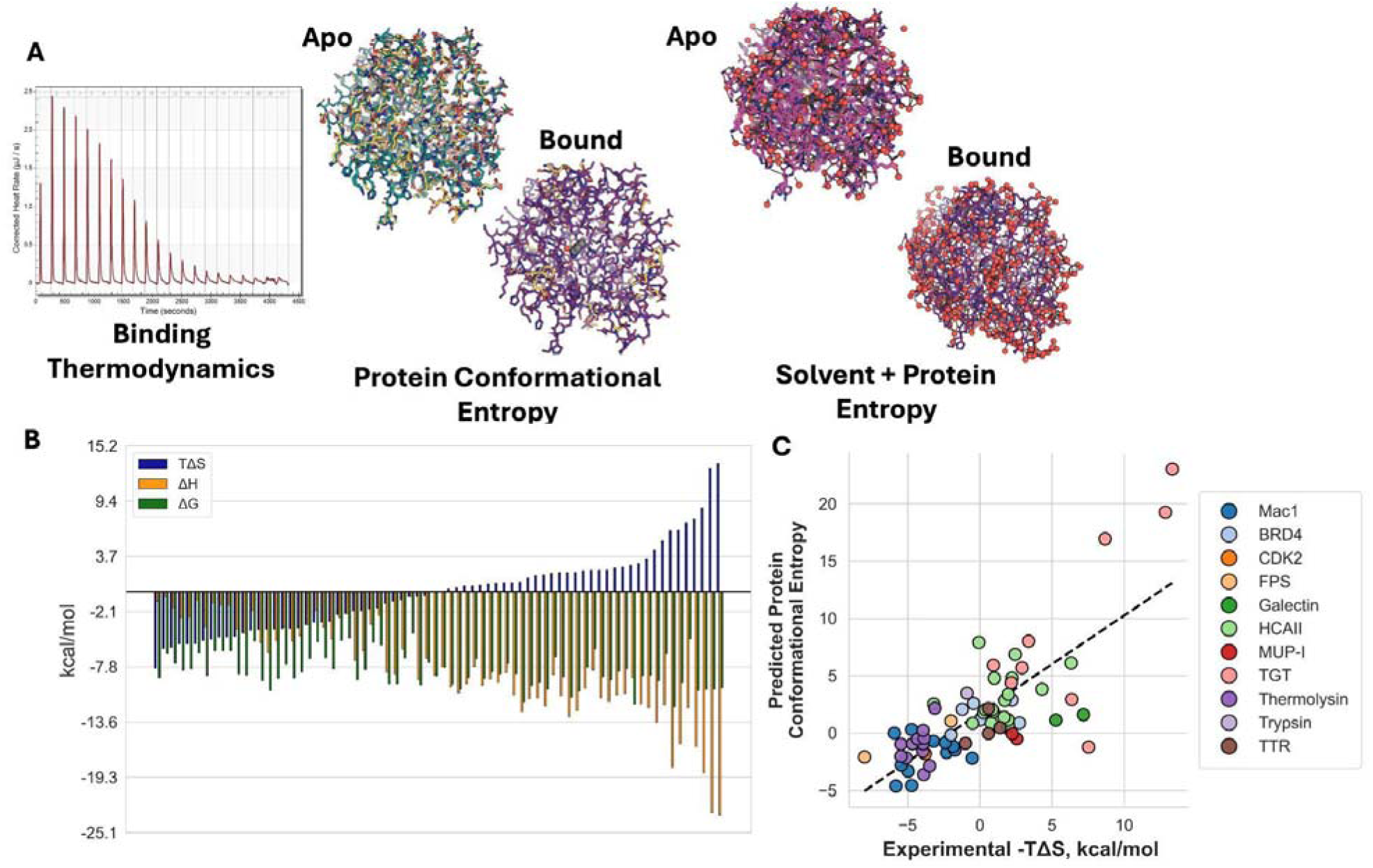
**A.** We obtained binding thermodynamic parameters by ITC from 14 proteins, comprising 72 protein-ligand pairs. Using multiconformer models, we estimated protein and solvent conformational entropies to identify correlations between ITC binding entropy and structural entropy estimates. **B.** Distribution of the thermodynamic binding contributions across all protein-ligand pairs ordered by entropy (n = 72). **C.** Correlation between protein conformational entropy as measured by crystallographic multiconformer structures and ITC-measured binding entropy. Each dot represents an individual protein-ligand complex, with each color representing a different protein (slope = 0.85, R² = 0.59). Higher binding entropy/protein conformational entropy indicates a more unfavorable binding entropy.

The strong observed correlation between protein conformational entropy and binding entropy, without explicitly accounting for solvent, likely arises for two reasons. First, we and others have observed that protein and solvent entropic contributions tend to covary [4,8] with structural rearrangements that alter protein conformational entropy, often simultaneously reorganizing interfacial and proximal water networks. Second, for the protein-ligand pairs studied here, protein conformational entropy may simply have a larger effect than solvent entropy [4].

We also noted that protein-ligand pairs from the same protein tended to cluster within a relatively narrow range of both structural and binding entropy values, with TGT being an exception[35,49]. We suspect that this limited dynamic range is driven by the ligand series used in each system, as we obtained ITC data from papers that often explore closely related ligands bound to the same protein, leading to similar binding thermodynamics. This pattern may also reflect intrinsic thermodynamic preferences of individual proteins, with some systems favoring more entropically driven binding mechanisms and others relying more on enthalpic stabilization. Previous work suggests this may be because rotational and translational entropy is constant for higher-affinity complexes, as residual rigid-body motion is limited for proteins compared to ligands within a protein-ligand complex [30]. Further binding-entropy analysis across a broader range of ligands is needed to test this hypothesis.

### Solvent Entropy Estimates are not Strongly Correlated with Binding Entropy

Having observed a strong linear correlation between protein entropy and binding entropy, we next investigated whether incorporating estimates of solvent entropy from crystallographic structures could further refine these predictions, as binding entropic effects arise from the collective behavior of protein, solvent, and ligand [50–52]. We first estimated changes in solvent-accessible surface area (SASA) between bound and unbound structures (**Methods**). Across all protein-ligand pairs, bound structures exhibited, on average, slightly greater solvent-accessible surface area (**Supplementary Figure 5)**. This increase was primarily driven by apolar SASA, which has been hypothesized to enhance solvent entropy by displacing ordered water molecules from apolar surfaces into the bulk solvent [53]. However, we observed no strong correlation between any SASA measure and binding entropy (total ΔSASA: R² = 0.0058; apolar ΔSASA: R² = 0.017; polar ΔSASA: R² = 0.0027; **Supplementary Figure 6**). Previous work has linked solvent-derived heat capacity estimates to entropy estimates using the relationship (ΔHeat Capacity = (−0.26)ΔSASAapolar + (0.45)ΔSASApolar) [7,54], prompting us to explore whether a similar relationship exists. However, we found no strong correlation between changes in heat capacity and binding entropy (R² = 0.066; **Figure 2A**).

**Figure 2.**
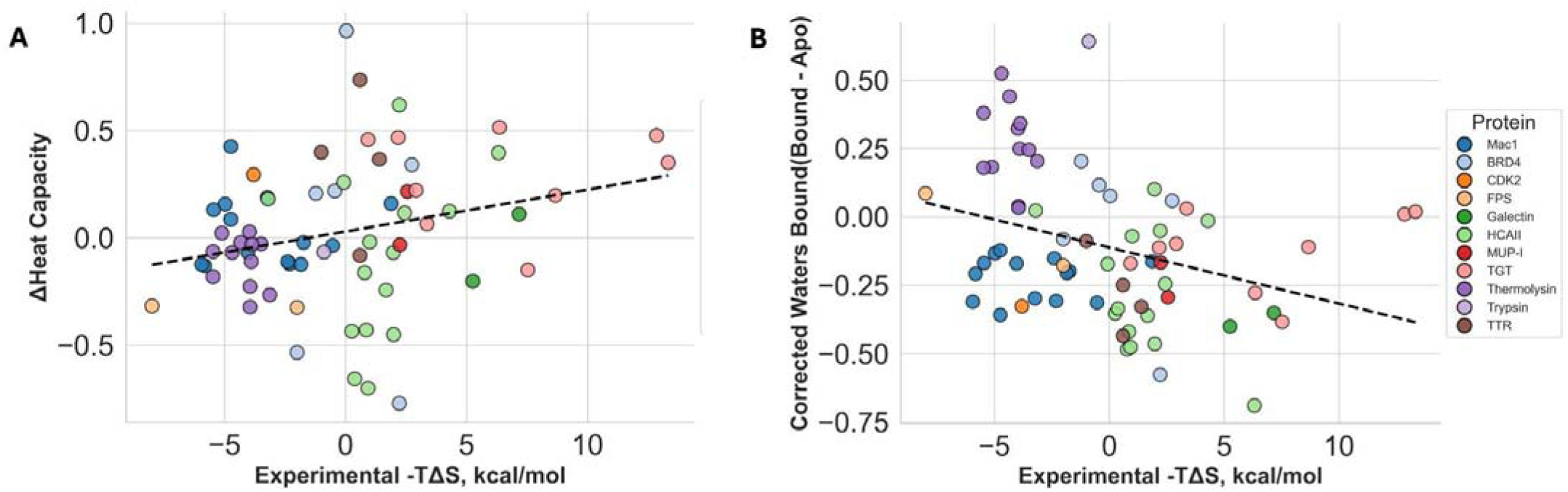
**A.** Estimates of differences in heat capacity with binding entropy (Slope = 0.020, R² = 0.066); **B.** Difference in the number of water molecules bound corrected for resolution and correlation with binding entropy (Slope = -0.021, R² = 0.11)

We reasoned that SASA measures alone may be insufficient, as they only indicate whether a residue is solvent-exposed, without capturing the complex thermodynamic behavior of water, particularly its hydrogen-bonding capacity or ‘iceberg’-like interactions at non-polar areas [50,51,55,56]. Therefore, we analyzed changes in the number of water molecules bound per residue, defined as water molecules within 3.2 Å of each residue (Methods). Across all protein-ligand pairs, most residues showed no change in bound water molecules (60.7%), while 22.8% gained and 16.5% lost water molecules (**Supplementary Figure 7A**). Notably, the mean difference in water molecule counts correlated with resolution differences between apo and bound structures, with the higher-resolution structure exhibiting more bound water molecules (R^2^ = 0.38; **Supplementary Figure 7B**). To address this confounding factor, we performed residual analysis to regress out resolution effects (**Methods; Supplementary Figure 8**). After correction, the distribution reversed: 56.7% of residues lost water molecules, while 43.3% gained them. We then tested whether differences in the number of water molecules, controlling for resolution differences, correlated with binding entropy. Although this slightly improved the correlation with binding entropy relative to heat-capacity estimates, it still had little power to explain binding entropy (**Figure 2B**; R² = 0.11).

### Binding Entropy can be Predicted from Protein and Solvent Entropy Estimates from Crystallographic Ensembles

We then tested whether solvent entropy estimates, combined with protein entropy, improved binding entropy prediction. We first used SASA values, including heat capacity; however, we did not observe any improvement over protein conformational entropy estimates alone (R² = 0.58; **Figure 3A; Supplementary Figure 9**). Finally, we evaluated whether changes in water molecule counts improved binding entropy predictions from crystallographic ensembles. Using a linear model, we found that resolution-corrected water counts improved prediction accuracy (R² = 0.69; -1.44 + 0.64 × Protein Conformational Entropy - 3.3 × Difference in Number of Waters (Corrected); **Figure 3B**). We observed slightly better improvements with uncorrected differences in water counts (R² = 0.704, **Supplementary Figure 10**). The consistent trends observed across proteins and ligands demonstrate that protein conformational entropy estimated from high-resolution crystallographic ensembles can provide semi-quantitative estimates of protein conformational entropy upon ligand binding.

**Figure 3.**
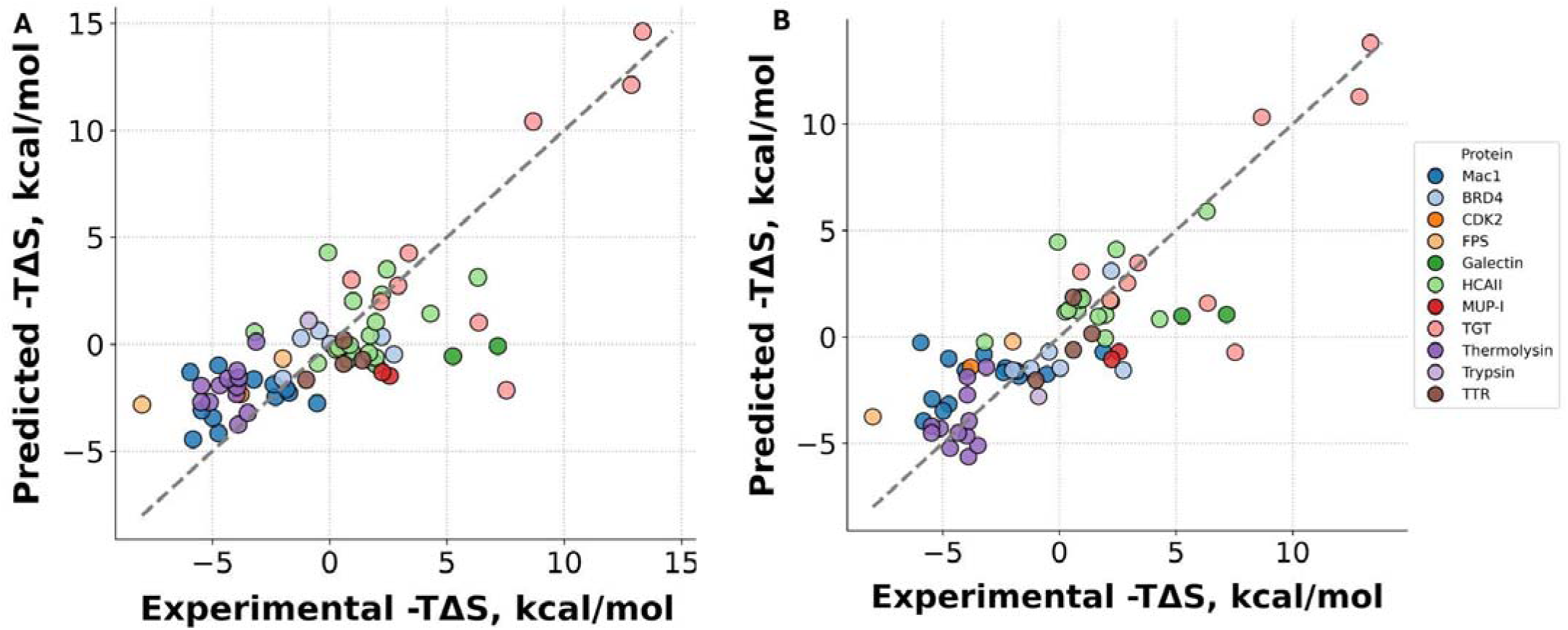
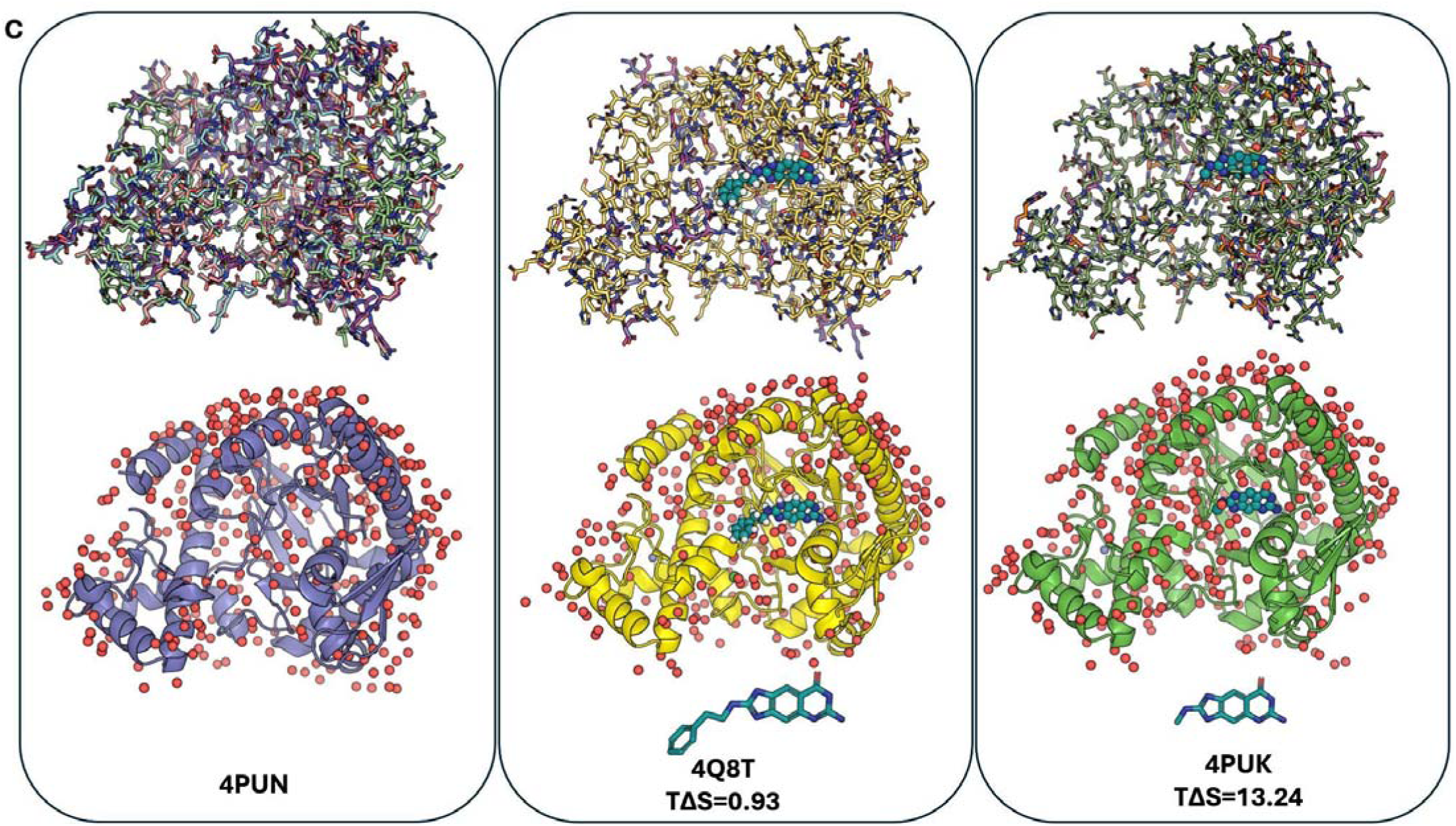
**A.** Correlation between predicted binding entropy values using estimates of heat capacity (R² = 0.59; TΔS = - 1.044 + 0.891*[Protein Entropy] + 1.097*[change in heat capacity]) **B.** Correlation between predicted binding entropy values using change in number of bound water molecules corrected for resolution (R² = 0.69; (TΔS = -1.45 + 0.64×[Protein Entropy] - 3.8×[Number of Water]) **C.** Protein conformational entropy and solvent differences between apo TGT (PDB: 4PUN) and two ligand-bound structures with dramatically different changes in binding entropy. Both ligands pay an entropic penalty, with 4Q8T paying a lower entropic penalty, derived from a smaller reduction in side chain alternative conformers (observed based on color differences of stick representations in the first row) and a smaller difference in the number of water molecules in the bound structure. 4PUK has more water molecules bound per residue with corrected resolution, further explaining the larger entropic penalty.

The most dramatic changes between binding entropies of the same protein were observed in TGT, with binding entropies ranging from 0.95 to 13.34 kcal/mol[35,49]. All entropies impose a penalty on free energy, though to varying degrees. We aimed to visually investigate what was driving these changes. In the apo state, nearly every residue exhibited alternative conformations, predominantly involving the backbone. Upon ligand binding, both complexes (4Q8T and 4PUK) showed substantial reductions in conformational heterogeneity, contributing to the entropic penalty of binding. Notably, 4PUK exhibited a larger estimated protein entropy change (11.31 vs. 5.91 kcal/mol·K) and a correspondingly larger experimental entropic penalty (13.34 kcal/mol). This complex also showed increased water occupancy (0.018 additional resolution-corrected waters per residue), whereas 4Q8T exhibited net water release (-0.17 fewer resolution-corrected waters per residue). These results demonstrate that both protein conformational restriction and differential solvent reorganization contribute to the distinct binding thermodynamics observed for these two ligands binding the same protein (**Figure 3C**).

### Hydrogen Bond Networks Show Moderate Correlation with Binding Entropy

We previously found that decreased residue packing and fewer protein–solvent hydrogen bonds in the bound state are associated with decreased binding entropy [8]. We aimed to assess whether these patterns generalize across different proteins. We excluded the Mac1 protein-ligand pairs from this analysis, as they were the source of our initial observations. To investigate the structural basis for entropic differences, we built a hydrogen bond network and computed graph-theoretic metrics that capture network connectivity, with nodes corresponding to hydrogen-bond-forming residues (**Methods**). We computed graph parameters, including node and edge counts. When examining protein-only features, we observed weak correlations between structural metrics and binding entropy, with the number of nodes (R² = 0.073) and edges (R² = 0.050) showing minimal predictive power (**Supplementary Figure 11**). Because local packing density restricts conformational sampling and correlates with residue-level entropy in folding studies [13], we tested whether packing metrics correlated with binding entropy. We quantified packing by counting neighboring residues within 5.0 Å of each residue’s heavy atoms. Packing density showed only a weak correlation with experimental binding entropy (R² = 0.13; **Supplementary Figure 12**), suggesting that the entropic penalties of binding are likely not driven primarily by static structural crowding.

However, incorporating water molecules into the hydrogen-bond network analysis (considering protein or water molecules as potential hydrogen-bonding nodes) revealed stronger correlations with binding entropy (nodes [R² = 0.38] and edges [R² = 0.24]; **Supplementary Figure 13**). This indicates that binding events that generate more extensive hydrogen-bond networks, encompassing both protein and water, tend to be less entropically driven. The notable improvement observed when water is included in hydrogen-bonding networks further supports the idea that protein and solvent jointly mediate the entropic cost of binding. These features may also help differentiate water patterns driven by distinct thermodynamic processes. For example, a more extensive hydrogen-bonding network involving solvent and protein would more strongly restrict solvent translation than a weaker network, thereby reducing translational entropy [57]. While the correlations remain moderate, the substantial increase in predictive power over protein-only metrics underscores the need to consider the entire protein-water system when analyzing binding thermodynamics.

## Discussion

We demonstrate that binding entropy can be predicted from crystallographic ensembles across diverse protein-ligand systems, enabling its quantification in structure-based drug design. We observe that accounting for both protein and solvent entropy is key to predicting this highly variable role in binding. This variability demonstrates that explicit entropy quantification is essential for effective design. Our approach addresses this gap by simultaneously optimizing both contributions using transferable metrics across globular proteins.

While entropy calculations require enumerating the partition function over all conformational states, they are computationally intractable due to high dimensionality and rugged energy landscapes. The crystallographic order parameter approach provides a practical alternative by measuring side-chain angular disorder as a proxy for conformational entropy [25,48]. Drawing on the Lipari-Szabo formalism [25], which relates order parameters to restricted angular motion, we capture local structural heterogeneity that reflects the system’s protein entropy. Further, our solvent estimations showed that using metrics based on explicitly placed solvent yielded a large improvement over implicit heat-capacity measurements [58]. Water molecules resolved in crystal structures represent those with sufficient occupancy and order to be detected, likely corresponding to waters that form stable, enthalpically favorable hydrogen bonds with the protein. The loss of water molecules upon ligand binding increases their rotational and translational entropy, contributing favorably to binding entropy. Conversely, the gain in ordered waters might stabilize specific conformational states, coupling solvent organization to protein entropy [51,59,60]. The improved binding-entropy predictions achieved by combining protein conformational entropy with water counts, compared with using either metric alone, indicate that these contributions are at least partially independent. However, we also found that changes in solvent-protein hydrogen-bonding networks contribute to binding entropy variability beyond what protein conformational changes alone explain. This finding suggests that protein and solvent entropy changes may reinforce each other [51,59].

The strong linear correlation with binding entropy validates our approach as an “entropy meter” for crystallography, extending NMR-based methods to the more accessible crystallographic domain [7]. However, several limitations warrant future improvement. Current metrics focus on single-residue angular motion rather than whole-side-chain descriptors and do not account for collective motions between residues or between protein and solvent. Improved protein conformational entropy metrics could be more directly grounded in statistical mechanics and could help illuminate correlations between protein and solvent entropy. Further, crystallographic ensemble models currently do not fully capture the conformational ensembles in the diffraction data, even before accounting for crystallographic artifacts. This limitation is particularly acute for water molecules, where resolution-dependent effects obscure the accurate quantification of solvent entropy [25,48,61]. Although our residual regression approach corrects for systematic resolution biases, it assumes uniform impact across all waters, an assumption that may fail if resolution differences correlate with genuine changes in structural heterogeneity rather than purely technical limitations. These challenges underscore the broader need to improve ensemble and solvent modeling methods. Finally, our dataset is dominated by closely related ligand series targeting the same proteins, potentially overweighting certain protein families and limiting the chemical diversity explored. Expanding this analysis to include more structurally diverse proteins and ligands would further test the generalizability of our findings.

Making entropy quantifiable from high-resolution crystallographic data enables thermodynamically balanced ligand optimization rather than enthalpy-focused design. We reveal that the dynamic ensemble, captured through multiconformer modeling, provides a practical means to quantify entropic contributions in structure-activity relationship campaigns, integrating this critical yet often overlooked free-energy component into rational drug design workflows. By developing an “entropy meter” based on crystallographic data, a widely available and accessible form of structural data, we hope to encourage equal consideration of enthalpic and entropic thermodynamic data in rational drug design workflows. This work bolsters an ongoing discussion in the field about how we extract, compare, and validate thermodynamic metrics in protein dynamics by proposing a new method based on crystallographic ensembles. While our dataset is relatively limited and increasing the diversity of protein-ligand complexes is important for future work, we believe this work can provide a foundation for predictive models to improve the efficacy of rational drug design pipelines. Our findings establish a foundation for entropically informed drug design, making the vision of designing ligands with optimized free-energy landscapes, rather than relying on enthalpic interactions alone, increasingly achievable.

## Methods

### Creation of multiconformer ensemble models

All structures were initially downloaded from PDBRedo[62] and were re-modeled using the qFit (version 2025.2)[29,47,63]. qFit structures were then processed through Phenix.refine (version 1.20)[47].

### Crystallographic Order Parameters

We computed crystallographic order parameter values for each residue (excluding glycine and proline) as a proxy for conformational entropy [48]. These values integrate both side-chain and backbone flexibility using two components: (1) the angle of alternative conformers (s2angle), calculated from χ1 dihedral distributions, and (2) atomic displacement data derived from normalized B-factors of alpha or beta carbons bonded to a hydrogen (s2ortho) [48]. The s2ortho term was corrected for resolution-dependent variation in B-factors using the following normalization:

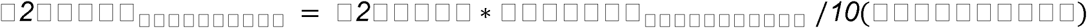

The final s2calc value was then obtained by combining both components multiplicatively:

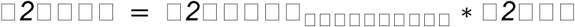

This combined metric captures both thermal motion and conformational variability, offering a robust estimate of per-residue rigidity. Values range from 0 (highly flexible) to 1 (fully rigid). We quantify side-chain rigidity using chi-1 rotamers because crystallographic density reports on side-chain conformation for all ordered residues, whereas NMR-derived order parameters are limited to methyl-bearing side chains and rely on inter-residue coupling<u>(Wand and Sharp 2018; Kasinath et al. 2013)</u>.

Protein conformational entropy was estimated as 0.25*number of residues*average crystallographic order parameter.

#### Solvent Accessible Surface Area

For each atom, we computed the SASA using FreeSasa [64]. We then divided the surface area into polar and apolar based on atom type. When calculating differences, we used continuous information

#### Water Molecule Analysis

We quantified local solvation by counting the number of water molecules within 3.2 Å of each residue in the structure. For each PDB file, we compared all non-hydrogen atoms of protein residues with the oxygen atoms of water molecules, and counted any water whose oxygen atom lay within 3.2 Å of any heavy atom in a given residue once. This analysis generated residue-level water contact counts, providing a quantitative measure of local hydration and solvent accessibility across the protein surface.

#### Water Molecule Resolution Correction

To remove systematic bias in water counts arising from differences in crystallographic resolution, we applied a resolution correction to all water count differences using linear residualization. We modeled the change in mean water count upon binding (bound minus apo) as a linear function of the resolution difference between paired structures. The expected contribution due solely to resolution was estimated by linear regression and subtracted from the observed water count change, yielding a resolution-corrected water count difference for each structure pair.

#### Hydrogen Bonds

Hydrogen bonds were identified if the donor–acceptor distance was ≤3.5 Å or the hydrogen–acceptor distance was ≤2.6 Å, and the donor–hydrogen–acceptor angle was≥120°. Networks were constructed as undirected graphs with residues as nodes and hydrogen bonds as edges, analyzed both with and without water molecules.

Network metrics included: basic properties (node/edge counts, density, degree distribution), connectivity (component analysis, clustering coefficients, transitivity), path metrics (average shortest path, diameter, radius), and centrality measures (betweenness, closeness, degree, eigenvector). Water-specific metrics quantified water-mediated versus direct protein–protein interactions. Per-residue metrics captured local connectivity and centrality within the hydrogen bonding network. We performed all analyses separately for networks that included and excluded water molecules to distinguish direct and water-mediated interactions.

#### Residue Packing

We quantified local packing density for each residue by counting neighboring residues within 5.0 Å using heavy-atom distances (excluding hydrogens and water). For each residue, we computed the number of neighboring residues, contact density (neighbors normalized by the number of heavy atoms in the residue), and average B-factor and occupancy values.

#### Ligand Properties

Ligand properties were obtained using RDKit via SDF files obtained from PDB structures.

## Data

All multiconformer modeled structures are in the Zenodo repository: https://zenodo.org/records/18209769

## Code

https://github.com/ExcitedStates/qfit-3.0

https://github.com/Wankowicz-Lab/ensemble_bioinformatic_toolkit

## Supplementary Figures

**Supplementary Figure 1.**
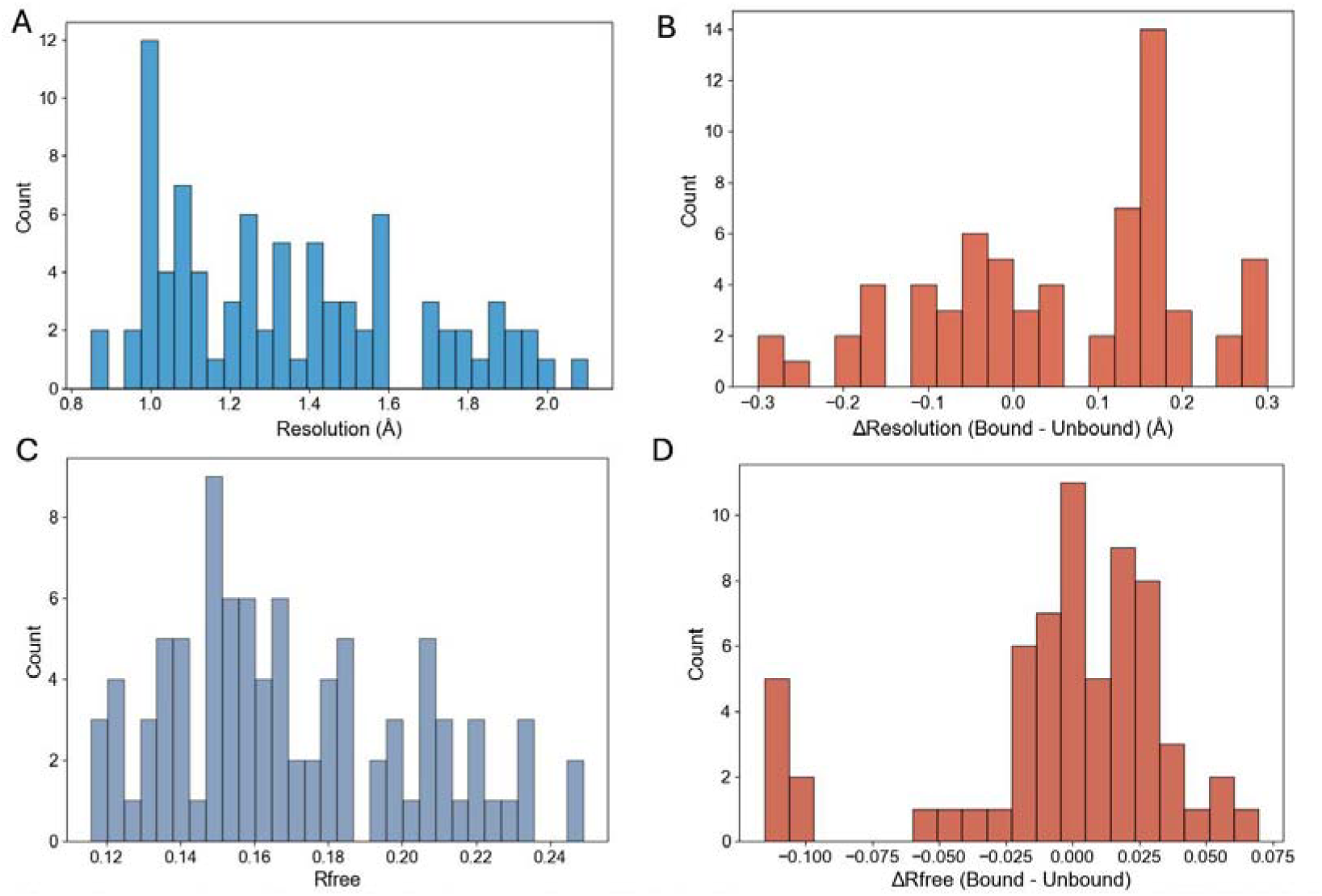
**A.** Resolution distribution across all PDBs (Median: 1.30 [IQR: 1.08-1.60]). **B.** Resolution differences between paired PDBs (Median: 0.06 [IQR: -0.06 - 0.15)). **C.** RFree distribution across all PDBs (Median: 0.16 [IQR: 0.15-0.19]). **D.** RFree differences between paired PDBs (Median: 0.010 [IQR: -0.010-0.027).

**Supplementary Figure 2.**
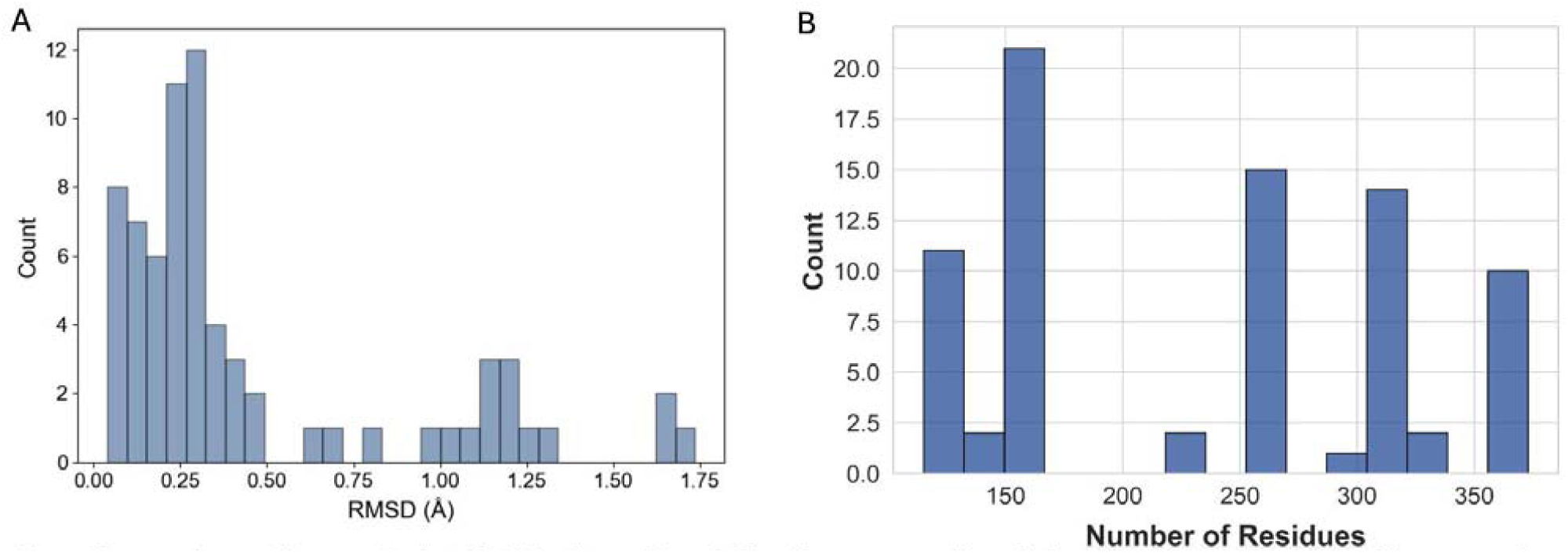
**A.** Distribution of protein size across the dataset by number of residue numbers. **B.** Alpha carbon RMSD between bound and unbound PDB pairs (Median: 0.27 [IQR: 0.19-0.46].

**Supplementary Figure 3.**
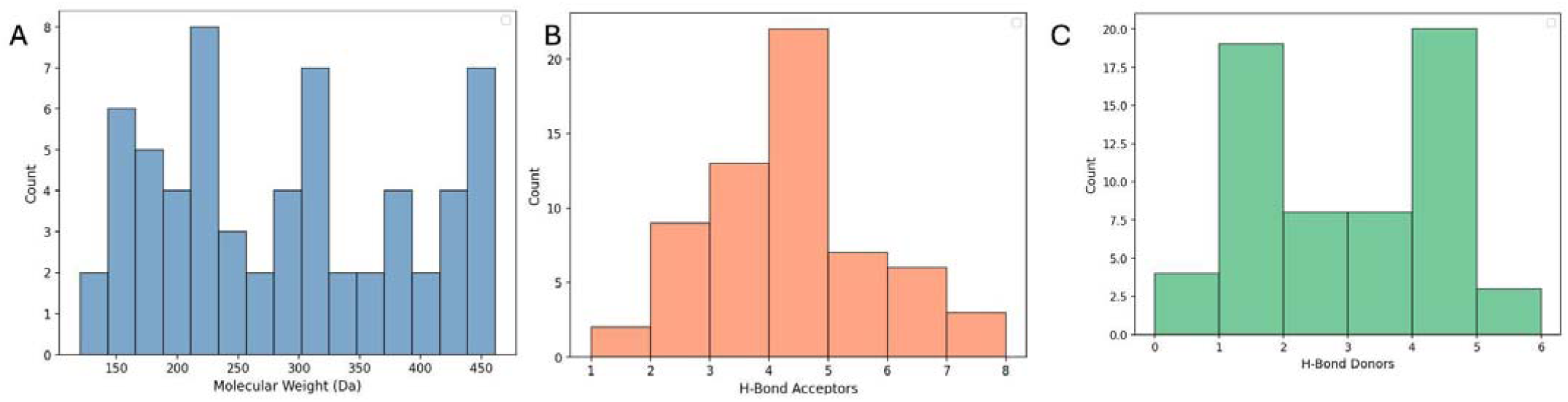
Ligand information. **A.** Molecular weight distribution across the dataset (Median: 284.68[102.46-461.50]) **B.** Hydrogen Bond Acceptor(Median: 4 [1-7]) C. Hydrogen Bond Donor (Median: 2 [0-5])

**Supplementary Figure 4.**
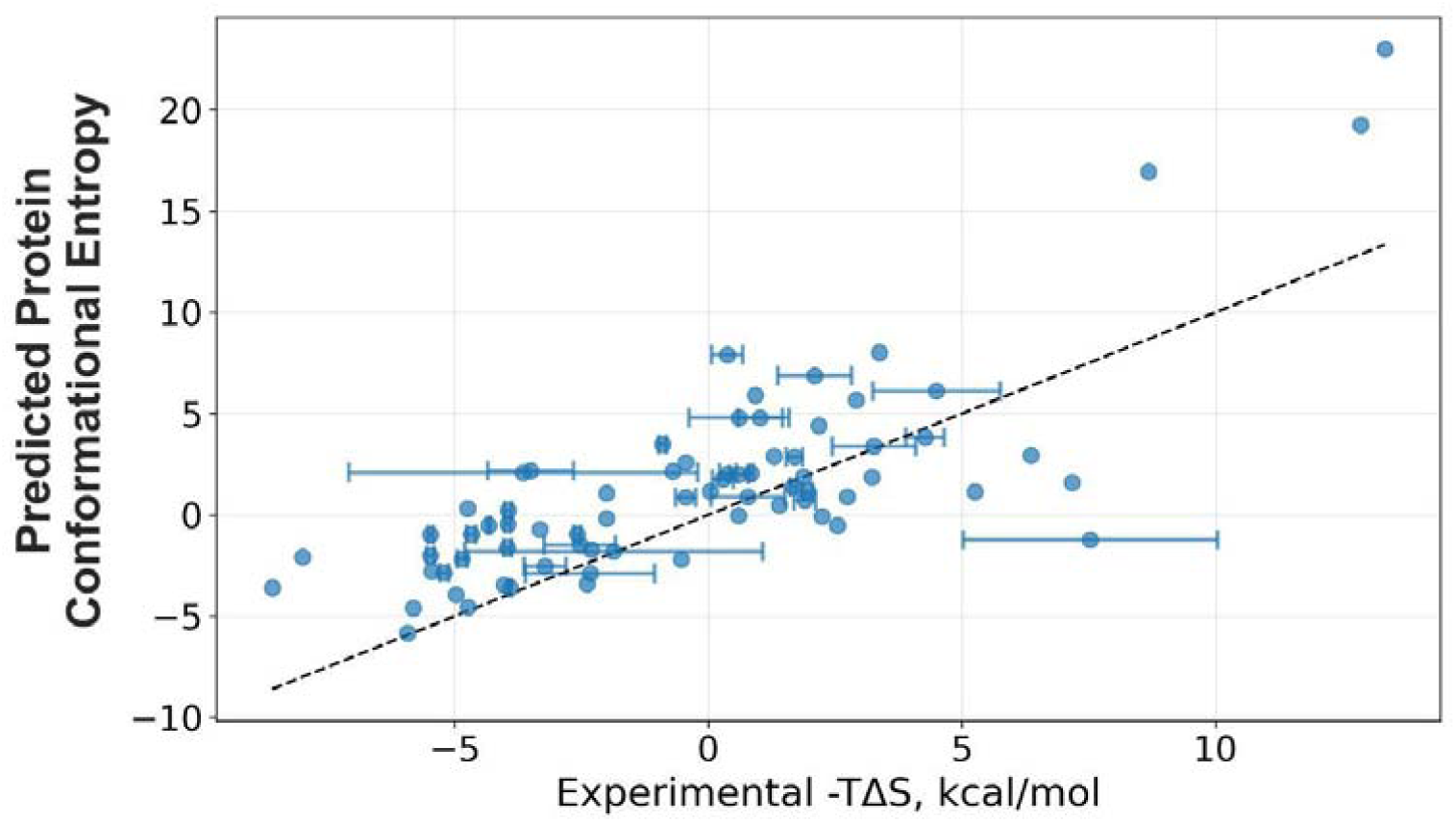
Correlation between ITC-measured binding entropy and protein conformational entropy metrics. Each point represents the mean binding entropy from ITC experiments, with error bars indicating the range of all measurements. Linear regression parameters: slope= 0.88, intercept= 1.63, R^2^ = 0.61.

**Supplementary Figure 5.**
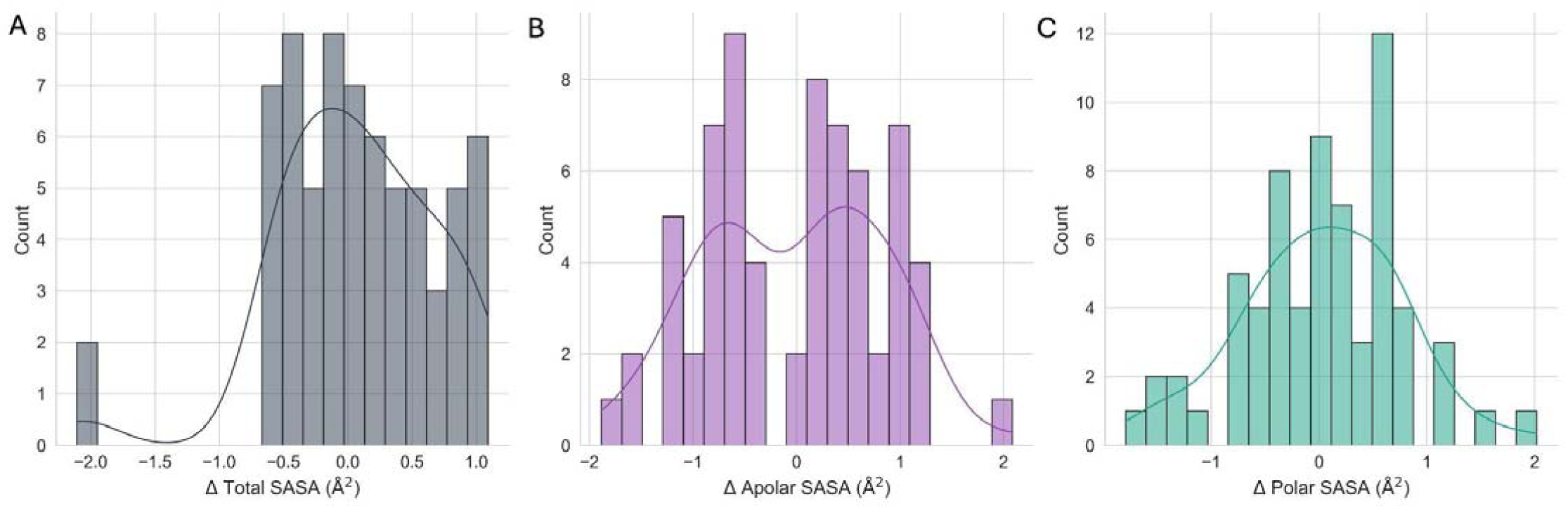
Distribution of change in SASA values across pairs. A. Total SASA changes (Median: 0.06, Mean: 0.05, Std: 0.62); B. Apolar SASA changes (Median: 0.16, Mean:-0.03, Std: 0.84); C. Polar SASA changes (Median: 0.07, Mean: 0.03, Std: 0.74)

**Supplementary Figure 6.**
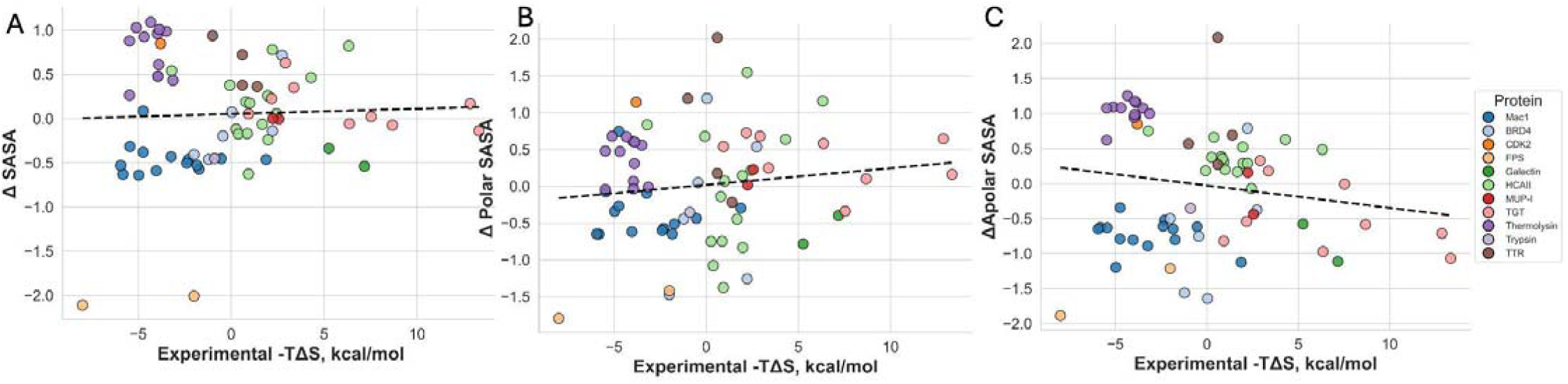
Correlation of SASA changes with binding entropy. A. Total ΔSASA: R^2^=0.0058; B. apolar ΔSASA: R^2^=0.0174; C. polar ΔSASA: R^2^=0.0027.

**Supplementary Figure 7.**
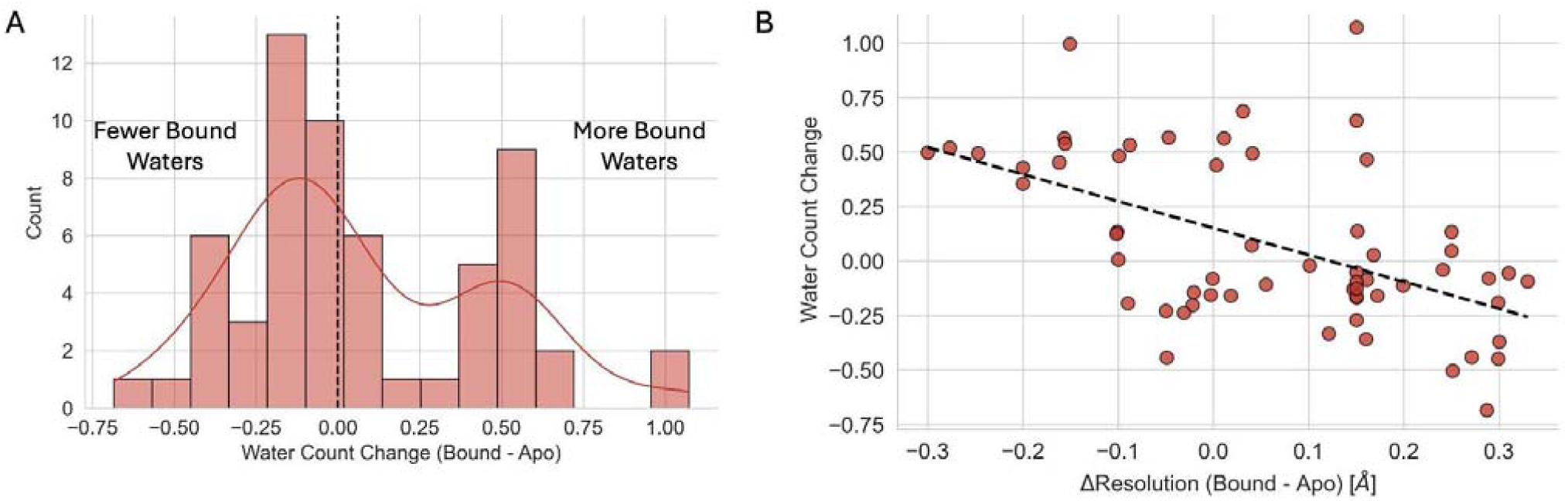
A. Distribution of average number of waters gained or lost between bound and unbound. B. Correlation between resolution differences and mean water count change per residue (R^2^=0.53, Slope=-1.232).

**Supplementary Figure 8.**
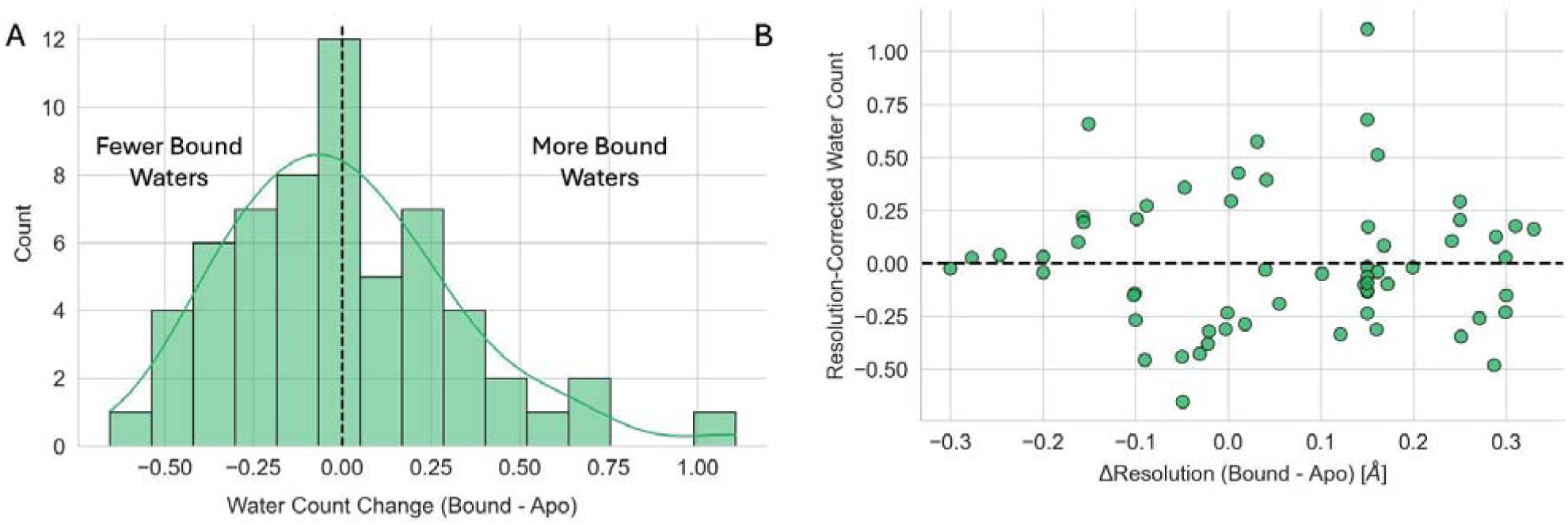
A. Distribution of average number of waters gained or lost between bound and unbound corrected for resolution differences between bound and apo. B. Correlation between resolution differences and mean water count change per residue (R^2^=0, Slope=0).

**Supplementary Figure 9.**
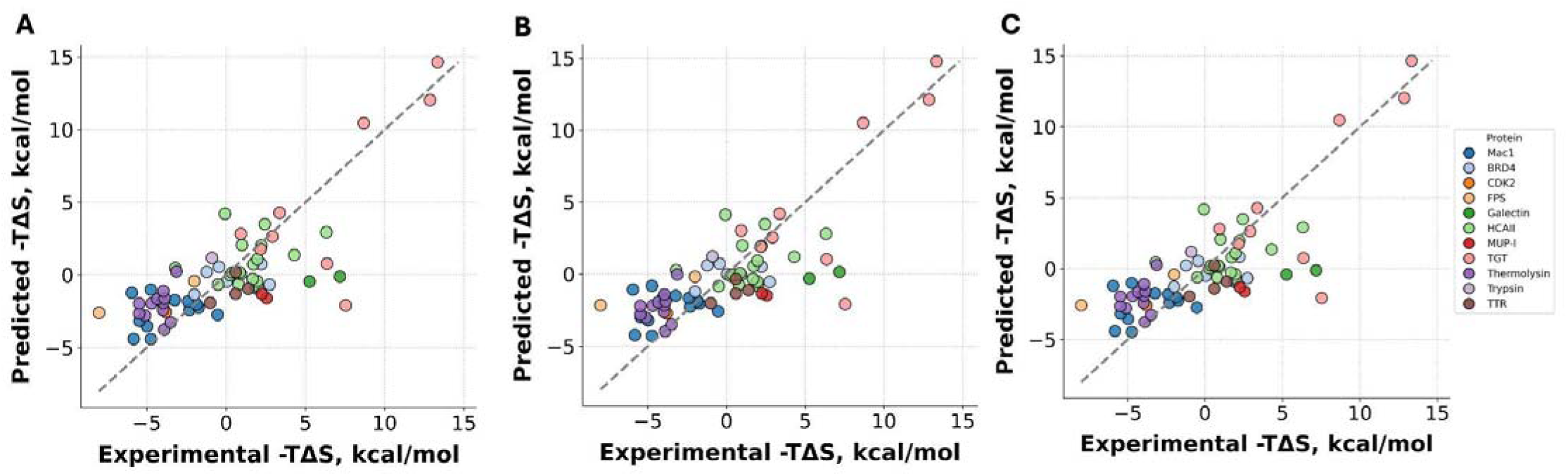
Contribution of solvent-accessible surface area components along with protein conformational entropy to experimental binding entropy. **A.** Model including total SASA (R^2^ = 0.588). **B.** Model including apolar SASA (R^2^ = 0.589). **C.** Model including polar SASA (R^2^ =0.587).

**Supplementary Figure 10.**
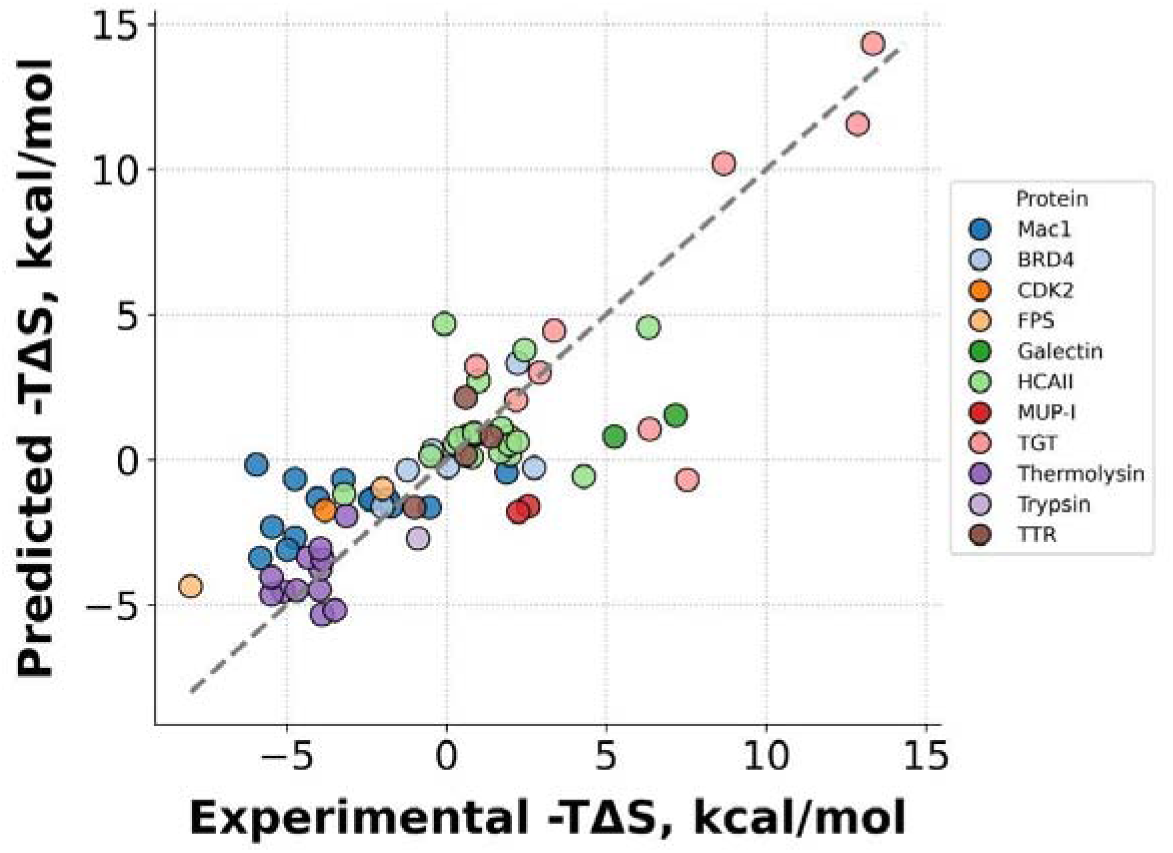
Linear model relating non-corrected protein conformational entropy and non-corrected water count to the entropic binding (R^2^ = 0.681)

**Supplementary Figure 11.**
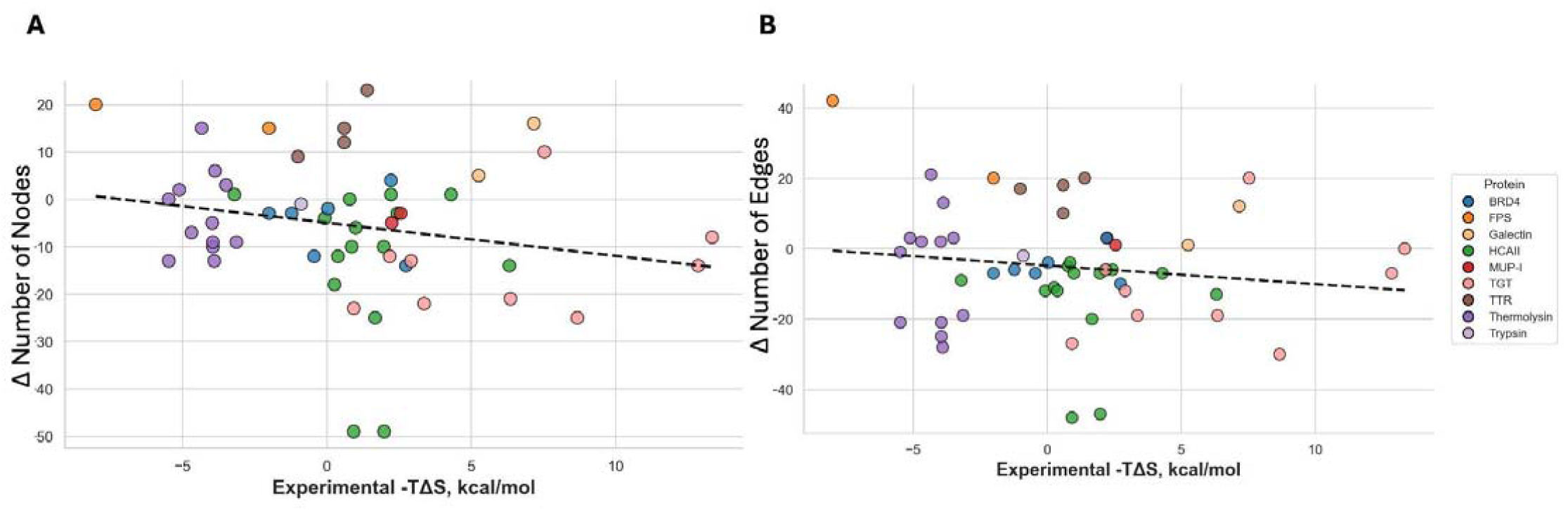
Hydrogen bond protein only. **A.** Correlation between the number of nodes and experimental entropy measurement (R^2^=0.0734) **B.** Correlation between the number of edges and experimental entropy measurement (R^2^=0.0489)

**Supplementary Figure 12.**
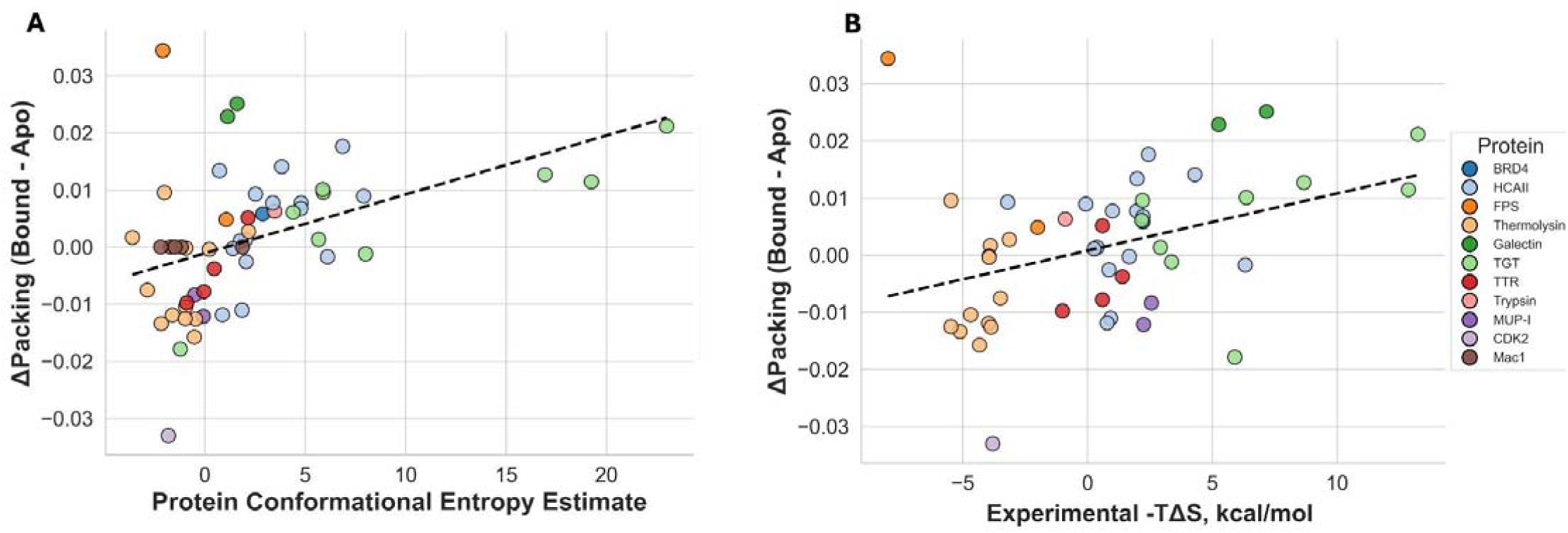
Packing and entropy measurement. **A.** Correlation between the difference in packing and protein conformational entropy estimates (R^2^=0.1976). **B.** Correlation between average number of waters gained or lost between bound and unbound with protein structural entropy estimates (R^2^=0.1318).

**Supplementary Figure 13.**
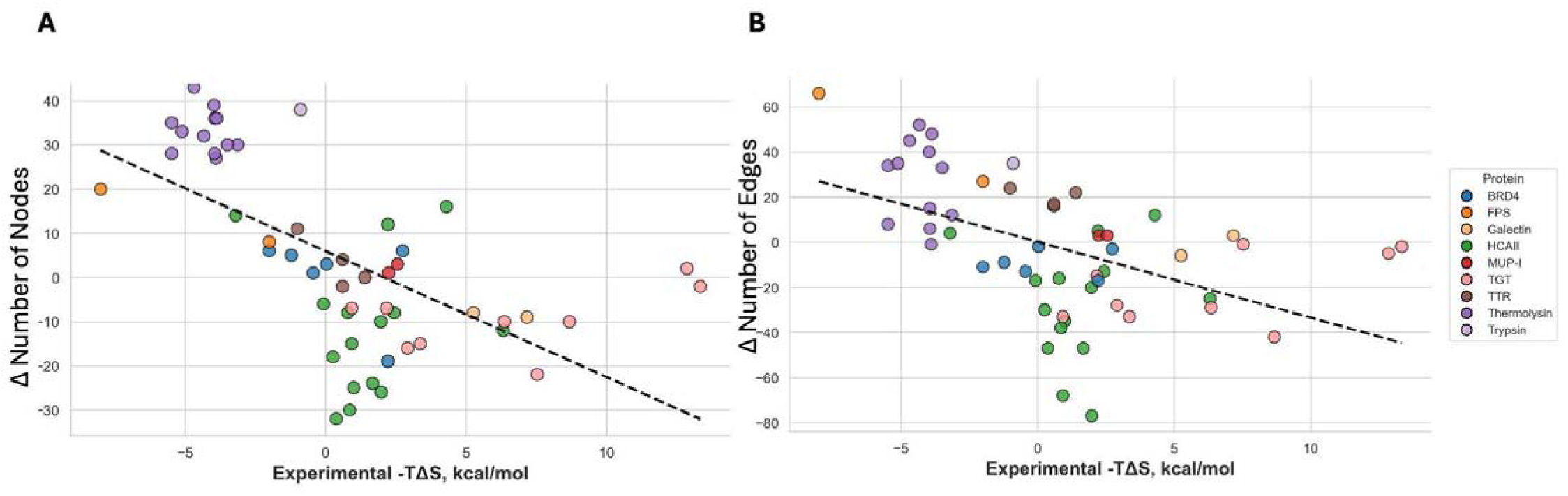
Hydrogen bond protein and solvent. **A.** Correlation between the number of nodes and experimental entropy measurement (R^2^=0.3765) **B.** Correlation between the number of edges and experimental entropy measurement (R^2^=0.2423)

## Notes

### Competing Interest Statement

The authors have declared no competing interest.

### Summary of Updates

This revision reflects responses to reviews within eLife. It includes additional details/rationalize around quantification we used throughout the manuscript.

### Funding

- American Cancer Society

